# Mast cells are not essential for pubertal mammary gland branching

**DOI:** 10.1101/2024.08.30.610382

**Authors:** Simran Kapoor, Jimmy Marsden, Clara M. Munz, Cyril Carvalho, Marlene Magalhaes Pinto, Bert Malengier-Devlies, Solvig Becker, Guillaume Seuzaret, Katelyn Patatsos, Ramazan Akyol, Marc Dalod, Amy Pedersen, Gillian Wilson, Rebecca Gentek

**Affiliations:** Institute for Regeneration and Repair, Centre for Reproductive Health, Centre for Inflammation Research, University of Edinburgh, UK; Institute of Medical Sciences, University of Aberdeen, UK & Research Institute for Environmental and Occupational Health, Université de Rennes I, France; Murdoch Children’s Research Institute, Royal Children’s Hospital Victoria, Australia; Aix-Marseille University, CNRS, INSERM, Centre d’Immunologie de Marseille-Luminy (CIML); Institute of Ecology and Evolution, School of Biological Sciences, University of Edinburgh, UK; Institute of Infection, Immunity and Inflammation, University of Glasgow, UK

**Keywords:** Mast cells, mammary gland, branching, puberty

## Abstract

Mast cells are long-lived, tissue-resident immune cells of the myeloid lineage with cardinal functions in allergy and atopic disease. They are now increasingly recognized also for protective roles e.g. against infections and venoms. Other functions originally assigned to mast cells in development and physiology, however, have been refuted, and for yet others, their true contribution remains uncertain. Mast cells have been implicated in promoting ductal branching in the pubertal mammary gland, the organ that produces and secretes milk in mammals, but these findings are based on mouse models that are not mast cell-specific. In this study, we therefore re-addressed the impact of mast cells on mammary gland branching using several complementary genetic models, including a new transgenic line. We report that neither constitutive deficiency of mast cells, nor their conditional ablation induced at puberty affected mammary gland branching. Our results thus dispute that mast cells promote this process in mice, at least in a unique and non-redundant manner. This study adds to a growing body of work clarifying the biological roles of mast cells, and further expands the toolbox available to the field of mast cell research.

## Introduction

The mammary gland is an organ specific to the mammalian reproductive system, and its primary function is milk production and secretion. It consists of milk ducts that spread throughout an adipose tissue pad and end in terminal end buds (TEBs), which facilitate duct branching and elongation during development and pregnancy. The mammary gland is also rich in resident immune cells and is indeed also of immunological importance: Transfer of immunoglobulin antibodies and immune cells through the milk can support offspring immunity (1), and the gland also provides an immunological barrier during suckling.

The mammary gland develops in a series of discrete steps ranging from embryonic to postnatal stages. In mice, mammary gland development is initiated prenatally at day 10.5-11 of embryonic development (E10.5-E11) with the formation of ectodermal disks that invade the underlying mesenchyme from E13.5 onwards and eventually form the nipples. The developing ductal epithelium then starts branching, resulting in a few rudimentary branches present at birth ((2), (3), (4), (5), (6)). These branches then remain largely quiescent and only grow allometrically until puberty, which marks the key period of mammary gland development. The start of ductal branching is often the first defining sign of puberty onset, and in humans is a developmental window of susceptibility to breast cancer ((7); (8)). During puberty, the previously rudimentary ducts grow and TEBs are formed. Cells at the end of the TEBs then proliferate to invade the fat pad, and branches form through bifurcation of TEBs (6). This process repeats until the ductal tree has invaded the entire fat pad. Throughout the reproductive cycle, the mammary gland undergoes further dynamic changes, enabling it to fulfil its physiological functions. With pregnancy, the secretory epithelium forms alveoli to prepare for lactation. Upon weaning, these alveoli undergo involution, and the gland returns to a state similar to that before pregnancy. Mammary gland remodeling is a carefully orchestrated process that relies on internal and external signals. Growth factors and hormones act on the mammary epithelium and stroma to facilitate branching and alveoli formation. Whilst fetal mammary gland development is independent of estrogen (9), it is suppressed by androgens in male mice. Testosterone exposure results in regression of mammary gland development (10), and androgens cause mesenchymal cells to condense around the developing epithelium, preventing further gland development ((10), (11)). On the contrary, pubertal branching depends on estrogen receptor α expression in epithelial cells (9). During puberty, progesterone acts on stromal cells to promote branching ((12), (2)). Finally, remodeling during pregnancy is induced by prolactin, which acts together with progesterone to promote alveolar growth. Local influences from stromal cells further contribute to remodeling. For example, fibroblasts in the mammary stroma communicate with the developing epithelium (13) by providing an extracellular matrix capable of maintaining the mammary glands (14).

Another player in the morphogenesis of mammary gland ducts are immune cells: Alveolarization during pregnancy is impaired in mice deficient in IL-4 and IL-13 (15), and pubertal duct elongation and branching are reduced in mice lacking eosinophils and macrophages in the mammary gland (16), (17), (18), (19)). This seems to also be true for mice deficient in mast cells, which show a reduction in the number of TEBs, as well as numbers and length of ducts during puberty (20). Mast cells are potent myeloid effector cells best known as mediators of allergy and anaphylaxis and for their implications in atopic disease. They also have protective roles, for example against bacteria and venoms ((21), (22), (23)) and by promoting avoidance behavior towards food allergens ((24), (25)). However, their contribution to normal tissue functioning and development remain heavily debated(26). This is at least in part due to limitations in the tools used to study mast cells. Their maturation and survival depend on stem cell factor, and inactivating mutations in its receptor Kit cause mast cell deficiencies. Traditionally, Kit-dependent mouse models have therefore been used to investigate mast cell functions (27). The usefulness of these models is limited, however, because Kit mutations also affect other immune and non-immune cells, as evidenced by several defects in Kit mutant mice that are unrelated to mast cells (27). This is of particular importance to the mammary gland, since Kit is expressed by the breast epithelium ((28)(29). Mast cells can be found near the ducts at the onset of puberty, and it has been suggested that they can directly interact with the ductal epithelium (20). Pubertal branching is stunted in the absence of mast cells, a defect that is rescued by adulthood through an unknown compensatory mechanism (20). However, these findings were made in mutant *Kit*^Wsh^ mice. It is currently unknown if mice with Kit-independent mast cell deficiency copy this phenotype, and hence, whether this defect is indeed attributable to the absence of mast cells.

In this study, we re-addressed the involvement of mast cells in pubertal mammary gland branching using Kit-independent genetic mouse models of mast cell deficiency, including a new, conditional and inducible transgenic line. Unexpectedly, we found no defects in mammary gland branching. Our data therefore dispute a critical contribution of mast cells to this developmental process, at least in a unique and non-redundant manner.

## Material and Methods

### Mouse lines

Rosa26^lsl-DTA^ (“DTA176”) mice were purchased from The Jackson Laboratory (strain #010527). *Mcpt5*-Cre mice were kindly provided by Axel Roers (Heidelberg, Germany), *Karma*^Cre^ mice (30) were gifted by Marc Dalod (Marseille, France), and the *Ms4a2*^hDTR^ line (also known as “red mast cell and basophil” or RMB mice (31), was obtained from Pierre Launay. *Ms4a2*^lsl-hDTR^ mice (official name: C57BL/6NRj-*Ms4a2*^tm2Ciphe^) are a new line we generated at the Centre d’immunophénomique (CIPHE) in Marseille (France). In these mice, the 3’-UTR of the *Ms4a2* gene encoding the FcεRI β chain includes a cassette composed of an internal ribosomal entry site, a sequence coding for the fluorescent protein tdTomato (tdT), a 2A cleavage sequence, and the human diphtheria toxin receptor (hDTR). In this line, expression of the hDTR and tdT transgenes is prevented by a stop codon that is flanked by loxP Cre recognition sites. This stop codon is removed upon Cre-mediated recombination, making these mice a Cre-inducible version of the *Ms4a2*^hDTR^ line.

### Animal ethics and housing conditions

All animal work was performed under project license (PPL) number PP1871024 in accordance with Animals (Scientific Procedures) Act 1986 (ASPA). Mice were bred and housed at the animal facility in the University of Edinburgh (Edinburgh, UK) under specific pathogen-free conditions at a controlled temperature of 22°C with a 12-hour light/dark cycle. Access to food (irradiated chow pellets) and water (reverse osmosis water) was available *ad libitum*.

### Diphtheria toxin-mediated mast cell depletion

Diphtheria toxin (DTx) (322326, Merck) was aliquoted in filtered PBS and stored at −80°C. *Ms4a2*^hDTR^ and *Karma*^Cre^:*Ms4a2*^lsl-hDTR^ mice were injected under general isoflurane anaesthesia at 4 weeks of age with 1µg DTx through the nipple into the fat pad of the 4^th^ mammary gland.

### Mammary gland harvest

The 4^th^ abdominal mammary gland fat pads were removed at the indicated ages for analysis by flow cytometry and/or imaging. The lymph node was excluded from the gland collected for cell suspension preparation and flow cytometry, and glands intended for imaging were left intact.

### Estrus staging

Were assessed, estrus staging performed by vaginal smear using 1x phosphate-buffered saline (PBS) onto Superfrost™ Plus Adhesion Microscope Slides (Epredia). Vaginal smears were air-dried, and then fixed in ice-cold 100% methanol, followed by staining with haematoxylin and eosin.

### Peritoneal lavage

Where done, peritoneal lavage was performed by injecting RPMI (supplemented with 2% FCS) into the abdominal cavity of mice immediately after cull, followed by gentle massaging to dislodge cells, and retrieval of the cell suspension using a syringe. The retrieved cell suspension was filtered through a 100µm filter membrane (John Stanair & Co) and stained for flow cytometry as detailed below.

### Preparation of single cell suspensions and flow cytometry

The dissected abdominal mammary gland fat pads were collected in RPMI supplemented with 2% FCS. Samples were cut manually into small pieces, followed by digestion in 0.8mg/ml Dispase II (Merck), 5mM CaCl_2_, 10mM HEPES (Merck), 0.1mg/ml DNaseI (Roche), 3mg/ml Collagenase D (Merck) in RPMI (supplemented with 2% FCS) at 37°C at 900rpm in a Thermo-Shaker (Grant-bio) for 20 minutes or until digested. Samples were filtered through a 100µm filter membrane (John Stanair & Co), and washed with FACS buffer (2% FCS, 2 mM EDTA in PBS) to obtain a homogenous cell suspension.

Cells were transferred to a 96 well-plate and incubated with 1:800 Zombie Live/Dead NIR (ThermoFisher Scientific) in PBS for 20-30 minutes at room temperature in the dark. Cells were then washed with FACS buffer and centrifuged at 530g for 5 minutes at 4°C. The supernatant was removed, and cells were incubated with 0.5% anti-mouse CD16/32 Trustain Fx (Biolegend), 5% mouse serum (Invitrogen) and 5% rat serum (Merck) in FACS buffer for 20-30 minutes at 4°C. Cells were then washed with FACS buffer and centrifuged at 530g for 5 minutes at 4°C. Cells were then washed with FACS buffer and centrifuged at 530g for 5 minutes at 4°C. Cells were then labelled with antibody mix diluted in FACS buffer and Brilliant Stain Buffer (BD) as detailed below for 20-30 minutes at 4°C.

| Marker | Fluorophore | Supplier | Clone number | Dilution |
| --- | --- | --- | --- | --- |
| CD45 | BUV395 | BD Biosciences | 30-F11 | 1:800 |
| CD117 (Kit) | BV650 | Biolegend | 2B8 | 1:300 |
| CD117 (Kit) | BV605 | Biolegend | ACK2 | 1:300 |
| CD11b | BV510 | Biolegend | M1/70 | 1:400 |
| CD64 | BV711 | Biolegend | X54-5/71 | 1:300 |
| F4/80 | BV786 | Biolegend | BM8 | 1:300 |
| CD63 | Pe/Cyanine7 | Biolegend | NVG-2 | 1:300 |
| ST2 | APC | Biolegend | DIH9 | 1:300 |
| ST2 | R718 | BD | U29-93 | 1:300 |

Cells were then fixed in 4% paraformaldehyde solution (PFA) for 15 minutes at room temperature away from light. Fixed cells were subsequently stained intracellularly with 1:10000 Avidin-AF488 (ThermoFisher Scientific) diluted in immunohistochemistry (IHC) buffer (2% Triton X-100) and 0.05 g/ml milk powder) for 1 hour at room temperature or overnight at 4°C in the dark. Cells were washed with FACS buffer and acquired in FACS buffer with counting beads (Invitrogen) (equivalent to 10.000 beads) added for quantification. Data was acquired using a 5L Fortessa (BD).

Cells were pre-gated as live, singlets. Mast cells were then defined as CD45^+^ CD11b^low^ F4/80^-^ CD64^-^ Kit^+^ ST2^+^ and/or Avidin^+^.

### Whole mount mammary gland imaging and morphology analysis

Dissected abdominal mammary gland fat pads were collected and spread onto Superfrost™ Plus Adhesion Microscope Slides (Epredia) and fixed for 24 hours in Carnoy’s buffer (75% ethanol, 25% glacial acetic acid). Fixed glands were then stained in Alum Carmine staining solution (0.1% carmine dye (ThermoFisher Scientific), 0.25% aluminium potassium sulfate dodecahydrate (ThermoFisher Scientific) in distilled water) for 2-7 days or until fully stained. Excess staining was removed by transferring the glands to a destaining solution (70% ethanol, 0.072% HCl) for 30 minutes or until excess staining was sufficiently removed. Glands were then dehydrated in an alcohol series (75%, 80%, 95%, 100% ethanol) and finally cleared in xylene overnight or until the fat pad was cleared. The slides were mounted in pertex mounting media and imaged. Images were acquired on a Leica Stereomicroscope at 0.8x magnification and analyzed using Fiji software. Branching parameters were measured from a line perpendicular to the length of the fat pad bisecting the lymph node. The distance covered by branches (in mm) was measured from the lymph node to the longest branch terminating in a terminal end bud. Numbers of TEBs and duct ends (TEDs) were counted in the direction of invading branching. Branching parameters were quantified by 2 independent parties blinded to experimental groups and/or genotypes.

### Statistical analysis

Unless otherwise stated, mean values and SEM as specified in the respective figure legends were calculated in Prism (GraphPad), and non-parametric t tests (Mann-Whitney). *P* values of less than 0.05 were considered statistically significant.

## Results

### Mast cells are present in postnatal mammary glands and appear activated at puberty

Like other immune cells, mast cells are commonly studied by flow cytometry, and they can also be identified in single cell transcriptomics datasets of samples obtained by tissue dissociation (32). However, to our knowledge, this approach has not exhaustively been applied to mast cells in the mammary gland. Therefore, we initially characterized mast cells in mammary glands from pubertal (5.5 weeks) and adult (aged 8-15 weeks) virgin mice by flow cytometry. Mast cells were identified as hematopoietic (CD45^+^) cells that lack markers for other myeloid lineages (CD64^-^ and/or F4/80^-^, CD11b^low^) and express Kit and the IL33-receptor alpha (ST2^+^) (Figure 1A). A characteristic feature of mast cells are their intracellular granules. These are densely packed with potent effectors and differ in their composition between distinct classes of mast cells. Mammary gland mast cells are of the connective tissue type ((32), (20)). These types of mast cells feature heparin-containing intracellular granules, which can be stained for with Avidin ((33), (34), (35)). Indeed, most Kit^+^ ST2^+^ mast cells also stained positive with fluorescently labelled Avidin in the mammary gland at puberty and in adulthood (Figure 1A). We thus used this as the primary gating strategy to identify mammary gland mast cells throughout this study.

**Figure 1:**
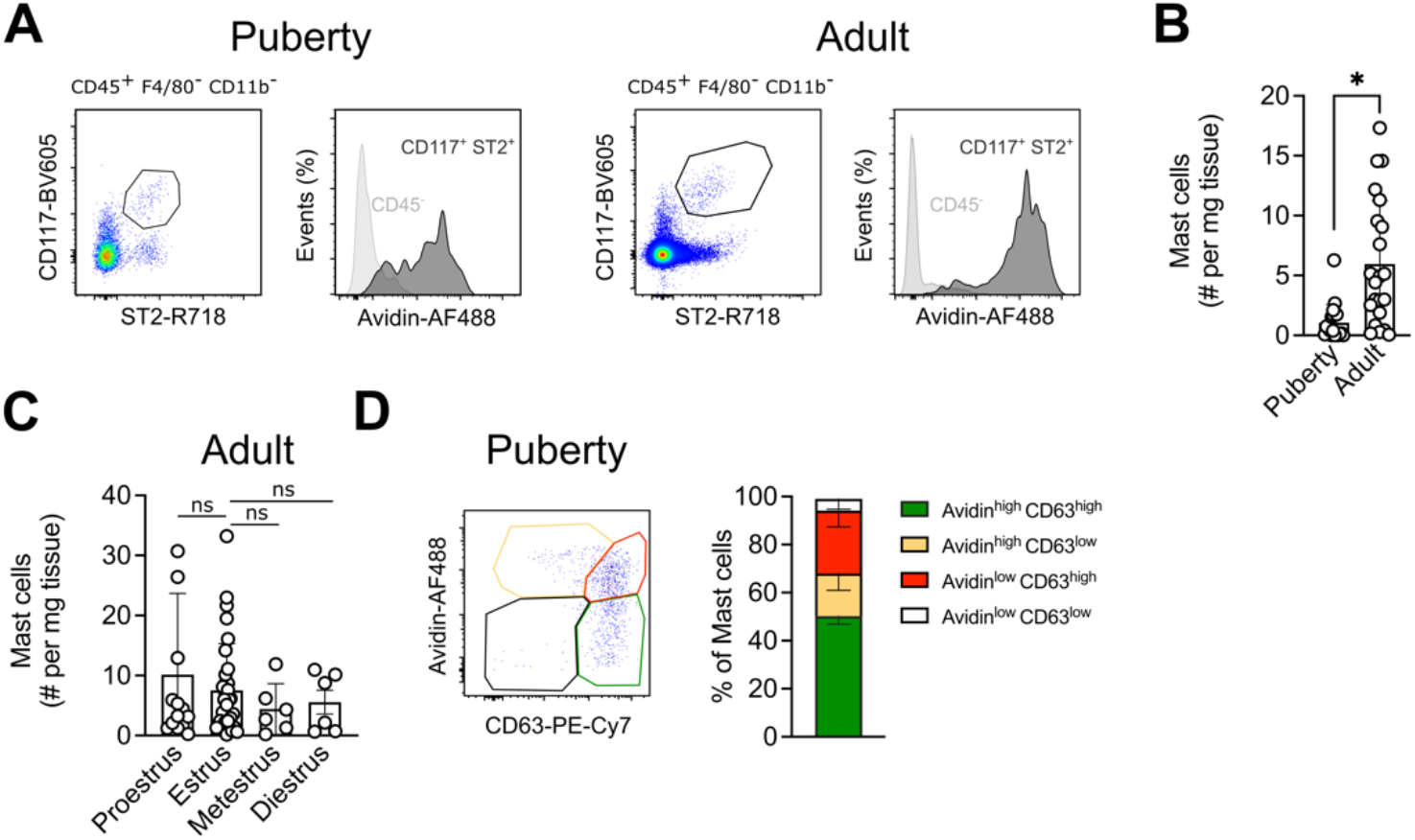
Mast cells are present in the postnatal mammary gland and appear predominantly activated at puberty. **(A)** Flow cytometry identification strategy for mast cells within the (left) pubertal (5.5 weeks) and (right) adult (8-15 weeks) mammary gland. **(B)** Quantification of mast cells (Kit^+^ Avidin^+^) in pubertal and adult mammary glands. Data in (A, B) are from at least 19 mice per age. **(C)** Mast cell numbers (Kit^+^ Avidin^+^) in adult mammary glands stratified by estrus stage. Data from minimally 6 mice per estrus stage. **(D)** Assessment of mammary gland mast cell activation states at puberty (5.5 weeks). Mast cells were gated as Kit^+^ ST2^+^ as shown in (A). Mast cells were left unstimulated, and the levels of degranulation were estimated by intracellular Avidin staining for heparin containing granules and surface staining of CD63. Representative (left) and quantified data (right) from 6 individual mice. Quantified data are shown as mean with error bars indicating the SD. \**P* < 0.05 as determined by Mann-Whitney test. ns = not significant.

Compared to puberty, mast cells were approximately 6-fold more abundant in adult glands (Figure 1B). We noted considerable variation between individual adult mice. In other organs of the reproductive system like the uterus, mast cell numbers vary with the estrus cycle ((36), (37)). Indeed, in rats, mast cells are more abundant at diestrus ((38)). Moreover, histamine levels are higher in the murine gland at estrus (((39); (40))). However, when we stratified mast cell numbers in the mammary gland by estrus stage, we did not find significant differences (Figure 1C).

The mammary gland consists of ductal epithelium and stromal cells, including adipocytes. In principle, mast cells could regulate ductal branching via both, locally restricted mechanisms as well as through longer-range signaling, since granules can travel far upon release. Indeed, mast cells are located throughout the gland in association with the ducts as well as the adipose tissue stroma ((20)). However, when we analyzed abdominal fat pads from adult male mice by flow cytometry, we obtained significantly less mast cells than in their female counterparts (Supplementary Figure S1A). Since male mice only have a rudimentary ductal tree, this suggests that most of the mast cells identified by flow cytometry in the females are not located in the adipose tissue, and instead, are likely mostly duct-associated.

Since mast cells are thought to promote ductal branching during puberty, we next wanted to assess the activation status of the mast cells present at this stage. Mast cell activation results in granule release, which can be measured by surface staining for the tetraspanin CD63, a component of granule membranes that becomes externalized upon degranulation (41). To gauge if mast cells can be activated in the pubertal gland, we thus determined surface expression of CD63 alongside levels of intracellular heparin-containing granules on freshly isolated cells *ex vivo*, without experimental treatments to trigger degranulation (Figure 1D). Using this assay, most mast cells are either actively degranulating (Avidin^high^ CD63^high^, on average 50.2%) or have recently been activated (Avidin^low^ CD63^high^; 26.2%), whilst only minor fractions appear to be in a non-active (Avidin^high^ CD63^low^, 17.8%) or immature or refractory stage (Avidin^low^ CD63^low^; 5%). This indicated that mammary gland mast cells are indeed functional at puberty.

In summary, we confirmed that mast cells are present in the adult and pubertal mammary gland. Moreover, we provide evidence that at puberty, most mast cells in the gland appear to be activated.

### Mammary glands are constitutively deficient in mast cells of *Karma*^Cre^:Rosa26^lsl-DTA^ mice

The data implicating mast cells in ductal branching in the mammary gland are primarily based on *Kit*^Wsh^ mice (20). To investigate if the transient delay in branching observed in these mice is indeed attributable to mast cells, we sought to use mice in which mast cell-deficiency does not depend on mutations in *Kit*.

A commonly used model in which mast cells are constitutively ablated in a Cre recombinase-mediated fashion are *Mcpt5*-Cre:Rosa26^lsl-DTA^ mice (42). In this model, Cre activity results in removal of a stop cassette that otherwise prevents expression of the Diphtheria toxin alpha subunit (DTA), thereby inducing death specifically in cells expressing *Mcpt5*, the gene driving Cre recombinase. *Mcpt5* encodes for a mast cell specific protease, which is shared by connective tissue-type mast cells like those found in the mammary gland. We thus reasoned that *Mcpt5*-Cre:Rosa26^lsl-DTA^ mice would be a good model to re-evaluate the functions of mast cells in this organ. However, in our hands, adult *Mcpt5*-Cre:Rosa26^lsl-DTA^ mice are not mast cell-deficient in the mammary gland (Supplementary Figure S2A). We therefore turned our attention to alternative Cre drivers. The *Karma* gene (also known as *Gpr141b*) encodes an orphan G-protein-coupled receptor. *Karma*^Cre^ mice were originally designed to target dendritic cells (DCs), but they also target connective tissue mast cells in the skin (30), and *Karma*^Cre^:Rosa26^lsl-DTA^ mice are profoundly deficient in peritoneal cavity mast cells (Supplementary figure S2B). We thus assessed the level of mast cell depletion in the postnatal mammary gland of *Karma*^Cre^:Rosa26^lsl-DTA^ mice. Considering our focus on pubertal maturation, we analyzed the window around puberty with higher granularity. Encouragingly, *Karma*^Cre^ mediated virtually complete ablation of mast cells in the mammary gland at all stages analyzed, i.e. prior to (3 weeks), at the onset (5 weeks), during (5.5 weeks) and at conclusion (6.5 weeks) of puberty, as well as in mature adults (8-12 weeks) (Figure 2A).

**Figure 2:**
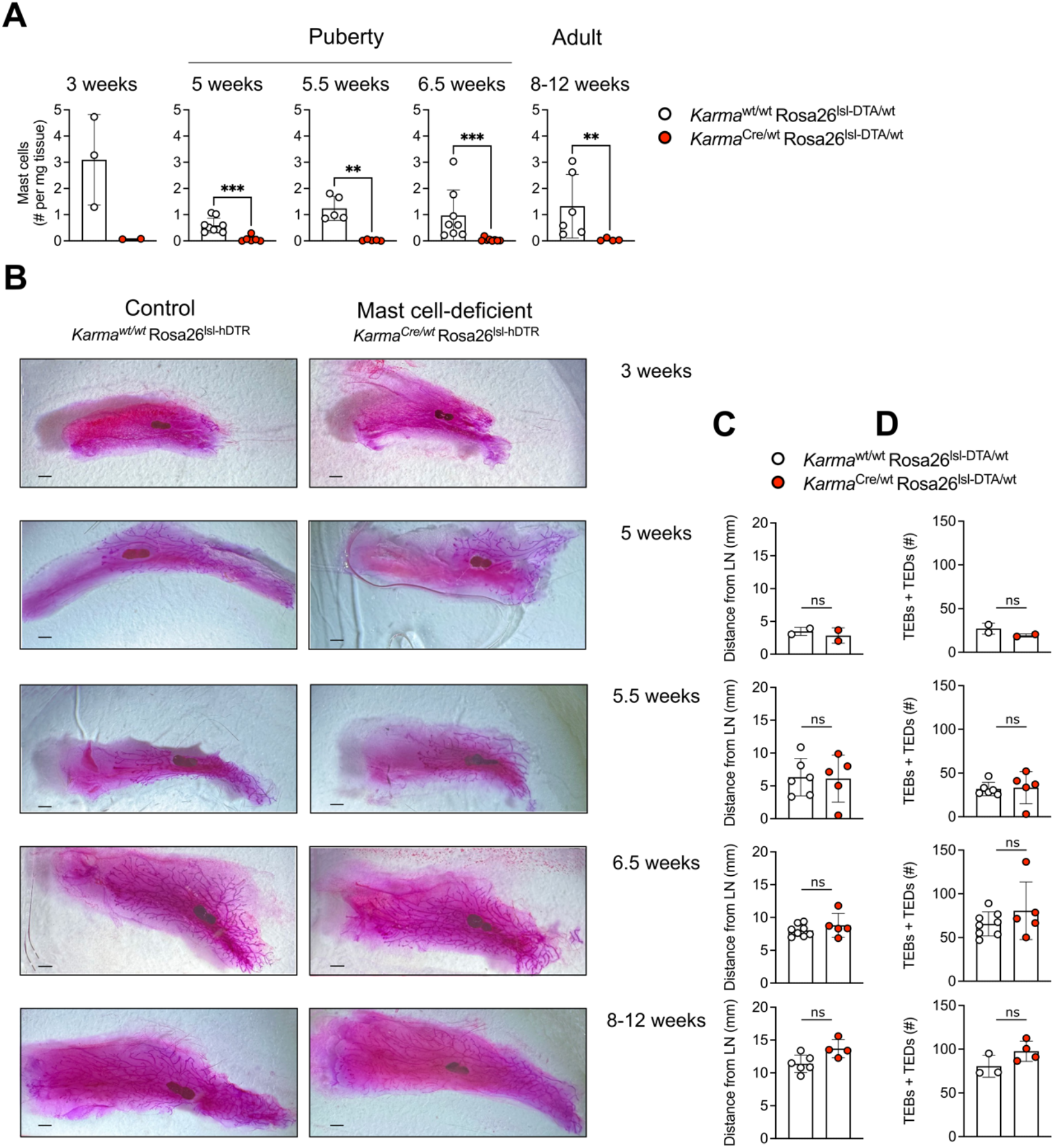
Constitutive mast cell deficiency has no impact on mammary gland branching. **(A)** Time course analysis showing mast cell numbers in *Karma*^Cre/wt^ Rosa26^lsl-DTA^ mice (red) compared to mast cell-proficient littermate controls (*Karma*^wt/wt^ Rosa26^lsl-DTA^ mice; white). Mast cells were quantified by flow cytometry at the indicated developmental time points. **(B, C, D)** Analysis of postnatal mammary gland branching on Carmine-stained mammary glands of mast cell-deficient (*Karma*^Cre/wt^ Rosa26^lsl-DTA^ mice; right/red) mice and mast cell-proficient littermates (*Karma*^wt/wt^ Rosa26^lsl-DTA^ mice; left/white) at the indicated ages. Scale bars = 1mm. The distance of the branching front (C) and the number of terminal end buds (TEBs) and duct ends (TEDs) (D) past the middle of the lymph node were measured. Data are representative (A) and cumulative (C, D) of at 56 individual mice from 10 independent experiments. Data in (A, C, D) are shown as mean with error bars indicating the SD. \*\*\**P* < 0.001, \*\**P* < 0.01 as determined by Mann-Whitney test. ns = not significant.

### Constitutive mast cell deficiency in *Karma*^Cre^:Rosa26^lsl-DTA^ mice has no effect on mammary gland branching

Having established *Karma*^Cre^:Rosa26^lsl-DTA^ mice as a tool for efficient mast cell deficiency in mammary glands, we next performed Carmine staining to investigate if these mice show any branching defects in the mammary gland. If mast cells did indeed promote pubertal branching, then these mice should exhibit impaired ductal branching like previously reported in *Kit*^Wsh^ mice (20). Because any defect might be transient in nature and compensated by adulthood, as is the case in *Kit*^Wsh^ mice (20), we performed a time course analysis. To our surprise, despite the absence of mast cells, *Karma*^Cre^:Rosa26^lsl-DTA^ mice did not show defects in branching at any stage of postnatal development (Figure 2B-D). Prior to puberty, at 3 weeks of age, mammary glands only contain rudimentary ducts. These were indistinguishable in mast cell-deficient *Karma*^Cre/wt^ Rosa26^lsl-DTA/wt^ and *Karma*^wt/wt^ Rosa26^lsl-DTA/wt^ control mice (Figure 2B). In our hands, the onset of puberty occurs at around 5 weeks of age, as evidenced by ductal outgrowth beyond the lymph node and the first signs of branching. Moreover, all mice used in our analyses during this time point were in proestrus/estrus stages. Between 5.5 and 6.5 weeks of age, ducts spread out and branches multiply, and the process is fully concluded by 8-12 weeks, when the entire fat pad is covered in ducts. However, at neither of these time points did we observe differences between mast cell-deficient *Karma*^Cre/wt^:Rosa26^lslDTA/wt^ animals and mast cell-proficient controls (*Karma*^wt/wt^ Rosa26^lsl-DTA/wt^. This was true for the extent of branching, as determined by the distance of the branching front beyond the lymph node (Figure 2C), as well as the number of TEBs and duct ends (TEDs) (Figure 2D). These data suggest that pubertal mammary gland branching occurs independently of mast cells, unlike currently thought.

### Specific ablation of mast cells at the onset of puberty has no effect on mammary gland branching in *Karma*^Cre^:*Ms4a2*^lsl-hDTR^ mice

A potential caveat of the *Karma*^Cre^:Rosa26^lsl-DTA^ model is that *Karma*^Cre^ also targets dendritic cells, in particular conventional (c)DC1 (30). DCs are present in the mammary gland, including subsets that resemble cDC1 (REFs: (43), (44)). DCs can be found already at 5 weeks of age (44) and may also be involved in the regulation of ductal branching, but in an inhibitory manner. Indeed, it has been reported that CD11c^+^ cells, which encompass DCs and macrophages, inhibit branching via a mechanism that involves antigen presentation to T cells (45). Moreover, cDC1 are reduced in *CD11c*-Cre:*Irf8*^fl/fl^ mice, and they show a trend towards moderately increased branching at 6 weeks of age (46). Whether these effects are due to targeting cDC1 has not been addressed. However, if branching was indeed promoted by mast cells and inhibited by cDC1s, then it is possible that simultaneous ablation of both has a net zero effect. As a consequence, any effects of mast cell deficiency might be “masked” by concomitant absence of cDC1s.

To circumvent this potential issue and exclude possible effects on other lineages, we revised an alternative genetic strategy that exploits the efficacy of *Karma*-Cre to enable selective ablation of mast cells, but not cDC1s or other lineages. To do so, we crossed *Karma*-Cre mice to a new transgenic line that we generated here. The latter is a derivation of the *Ms4a2*^hDTR^ model, also known as “Red mast cell and basophil” (RMB) mice (31), in which the human Diphtheria toxin receptor (hDTR) is expressed under control of the *Ms4a2* gene, which encodes the beta subunit of the IgE receptor. In these mice, mast cells and the closely related basophils are sensitive to Diphtheria toxin (DTx)-mediated ablation, because they both express *Ms4a2* (31). Reassuringly, delivery of the toxin into the mammary fat pad of pre-pubertal *Ms4a2*^hDTR^ mice resulted in complete loss of mast cells in the mammary gland and peritoneal cavity (Figure S3). To further refine this approach, we rendered the *Ms4a2*^hDTR^ system conditional by introducing a loxP-flanked stop cassette in front of the sequence coding for hDTR (see Material and Methods). This approach restricts DTx-sensitivity to cells that have undergone Cre-mediated recombination and express *Ms4a2*. We refer to this new line here as *Ms4a2*^lsl-hDTR^. Intercrossing *Karma*^Cre^ and *Ms4a2*^lsl-hDTR^ mice results in a combined model in which only mast cells are susceptible to depletion by DTx, but not e.g. cDC1s or basophils, since these respectively lack expression of *Ms4a2* or *Karma*.

Indeed, administration of a single dose of 1µg DTx by subcutaneous injection into the mammary fat pad just prior to the onset of puberty at 4 weeks resulted in the absence of mast cells in the mid-pubertal gland of *Karma*^Cre^:*Ms4a2*^lsl-hDTR^ mice at 5.5 weeks (Figure 3A), the time point when mammary gland remodeling was most active in our hands. We then used this strategy to determine if conditional ablation of mammary gland mast cells with the onset of puberty affects ductal branching in this critical window. Like in the *Karma*^Cre^:Rosa26^lsl-DTA^ model, however, we found no difference in the extent of duct migration into the fat pad and branching between mast cell-deficient (*Karma*^Cre^:*Ms4a2*^wt/hDTR^) and control mammary glands (*Karma*^Cre^:*Ms4a2*^wt/wt^) (Figure 3B, C).

**Figure 3:**
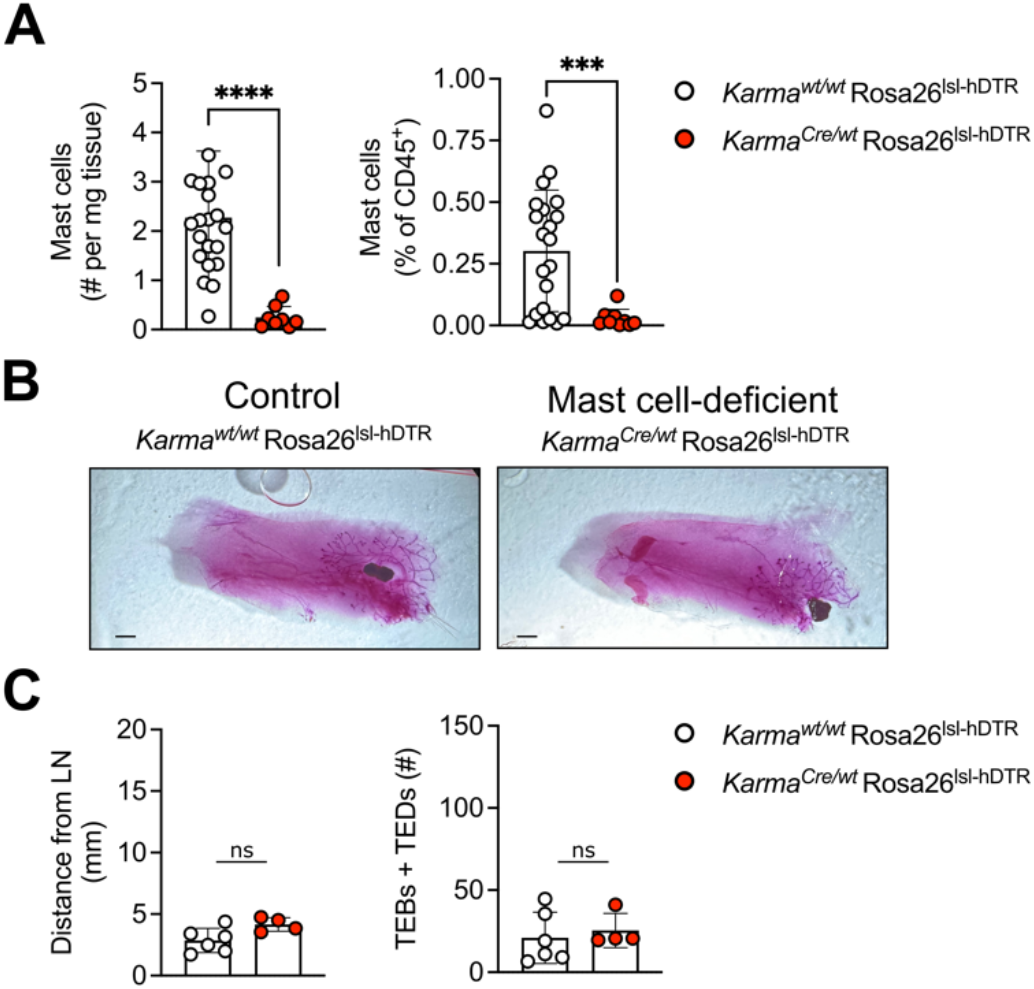
Mast cell deficiency induced at puberty does not affect mammary gland branching. To induce mast cell ablation, *Karma*^Cre/wt^ Rosa26^lsl-hDTR^ mice were treated with 1ug Diphtheria toxin at 4 weeks of age by subcutaneous injection into the mammary gland fat pad. Littermate control animals (*Karma*^wt/wt^ Rosa26^lsl-hDTR^) were treated the same way. Animals were analyzed at puberty (5.5 weeks of age). **(A)** Mast cell numbers and relative abundance within hematopoietic compartment as determined by flow cytometry. **(B, C)** Branching as determined by Carmine staining of pubertal mammary glands from 5.5 weeks-old mast cell-deficient (*Karma*^Cre/wt^ Rosa26^lsl-DTA^ mice; right/red) and mast cell-proficient mice (*Karma*^wt/wt^ Rosa26^lsl-DTA^ mice; left/white). Scale bars = 1mm. The extent of branching was measured as distance of branches and the number of terminal end buds (TEBs) and duct ends (TEDs) from the middle of the lymph node. Data are representative (B) and cumulative (A, C) of at 39 individual mice from 4 independent experiments. Data in (A, C) are presented as mean with error bars indicating the SD. \*\*\*\**P* < 0.0001, \*\*\**P* < 0.001, \*\**P* < 0.01 as determined by Mann-Whitney test. ns = not significant.

Collectively, our data challenge the notion that mast cells promote pubertal branching in the mammary gland, at least in a unique, essential manner.

## Discussion

In this study, we investigated if mast cells promote ductal branching of the mammary gland during puberty, as previously reported in *Kit*^Wsh^ mice (20). Using two complementary alternative genetic models for either constitutive or selective inducible ablation of mast cells, we found no evidence for an involvement of mast cells in this developmental process.

There are several possible explanations for the absence of a branching phenotype, which are not mutually exclusive. First, as introduced, expression of Kit is not restricted to mast cells, or even the immune system. This includes the epithelium of the mammary gland ((28), (29)). Indeed, many roles previously assigned to mast cells in *Kit* mutant mice have been refuted in Kit-independent models. For example, *Cpa3*^Cre^ or “Cre master” mice lack mast cells due to Cre-mediated cytotoxicity, and unlike *Kit* mutant mice, these are susceptible to antibody-induced arthritis (47), autoimmune encephalomyelitis (47) and diet-induced obesity (48). The latter has also been confirmed in *Mcpt5*-Cre Rosa26^lsl-DTA^ mice (49). In the mammary gland, Kit is expressed in epithelial cells, where it has been associated with growth and maintenance ((28), (50), (29), (51)). Our findings thus add to a growing body of work showcasing the limitations of Kit-based models for mast cell research, and for the first time, highlight these in a developmental context. Whilst the use of Kit mutant versus Kit-independent mice could explain the discrepancy between our findings and the literature, we acknowledge that Lilla and Werb (20) also observed stunted branching at puberty in mice treated with chromolyn sodium to prevent mast cell degranulation, as well as in mice deficient in Dipeptidyl peptidase I (DPPI) (20). Chromolyn sodium is commonly used as mast cell stabilizing agent, but can also impact additional cell types, including other granulocytes, lymphocytes, and stromal cells like fibroblasts, endothelial and epithelial cells. Whilst some of these may be indirect, direct effects of chromolyn sodium on other cell types have also been reported *in vitro* and *in vivo* in the absence of mast cells ((52), (53), (54)). In a similar manner, DPPI is needed for the activation of mast cell proteases, notably chymases (55). However, it also mediates activation of proteases in cytotoxic lymphocytes, monocytes and neutrophils (56).

Second, although we found no branching defects in either *Karma*^Cre^:Rosa26^lsl-DTA^ or *Karma*^Cre^:*Ms4a2*^lsl-hDTR^ mice, it remains possible that mast cells do contribute to the regulation of gland remodeling. However, rather than having unique, non-redundant effects, they may normally do so in cooperation with other cell types. Alternatively, other cells may functionally compensate at least when (only) mast cells are absent, even if they physiologically are not the main drivers of the process. Such concerted action of mast cells with other cells has been described in the uterus. To facilitate increased blood flow to growing fetuses, uterine spiral arteries need to be transformed from thick-walled vessels to vessels with enlarged lumen during gestation. Fetal growth restriction occurs if this process is insufficient. This occurs in *Kit*^Wsh^ mice and can be rescued by reconstitution with wild-type mast cells (36). On the contrary, *Cpa3*^Cre^ mice only show a mild spiral artery remodeling defect that does not cause fetal growth restriction (57). Spiral artery remodeling however requires smooth muscle cells to become apoptotic, and this is mediated by *Mcpt5* (58). In the uterus, Mcpt5 is produced not only by mast cells, but also a subset of natural killer (NK) cells (58). Akin to *Cpa3*^Cre^ mice that lack only mast but not NK cells, those lacking just NK cells also only show a mild defect in spiral artery remodeling (57). Profoundly defective uterine vascular remodeling and fetal growth restriction are therefore only observed in mice with combined deficiency in both, mast cells and NK cells, achieved either genetically in *Mcpt5*-Cre Rosa26^lsl-DTA^ mice or via antibody-mediated depletion of NK cells in mast cell-deficient *Cpa3*^Cre^ mice (58). Moreover, numbers of uterine mast cells are increased in IL-15 knockout mice which are genetically deficient in NK cells, and conversely, uterine NK cell numbers are higher in *Kit*^Wsh^ mice compared to mast cell-proficient controls (59). This underscores that a compensatory increase in other immune cells can indeed occur in the absence of mast cells, and that this may mask defects. Given how vital spiral artery remodeling is to reproductive success, it is perhaps unsurprising that a complex regulatory system with several layers has evolved. In the mammary gland, such a scenario could explain why branching is unaffected in mice with selective mast cell deficiency, i.e. *Karma*^Cre^ *Ms4a2*^lsl-hDTR^, but is delayed at puberty in DPPI knockout mice or following treatment with chromolyn sodium (20), which may also affect other cells, as discussed above.

Finally, it is also possible that diverse types of mast cells exist within the mammary gland, which may have distinct biological roles. Indeed, it is increasingly recognized that mast cells are heterogenous beyond the classic dichotomy of connective- and mucosal-type mast cells ((60), (61)). Mast cells also dynamically change across life stages ((62), (61), (63)). One aspect in which mast cells differ is their developmental origin. At mucosal sites, mast cells are predominantly derived from hematopoietic stem cells (HSC) ((61), (63), (64), (65)). In many connective tissues, however, mast cells are long-lived and self-maintain largely independently of HSCs in the bone marrow ((61), (63), (64)). They also retain sizeable populations of the first mast cells that originate from yolk sac erythro-myeloid progenitors ((63), (64)). Mast cells from distinct sources also retain functional differences, at least upon immunological challenge e.g. by infections, or in dynamic settings like tissue remodeling during development (66), Any such heterogeneity has not yet been resolved for mammary gland mast cells, but it is conceivable that they comprise subpopulations which e.g. are more or less responsive to hormones, and may thereby differently impact branching, perhaps even in opposing ways. Since *Karma*^Cre^ Rosa26^lslDTA^ and *Karma*^Cre^ *Ms4a2*^lsl-hDTR^ mice lack all mast cells in the mammary gland, this is a possibility that we cannot account for here.

### Data Limitations and Perspectives

Our primary aim here was to re-evaluate in Kit-independent mouse models if mast cells are essential for mammary gland remodeling at puberty. As such, one limitation is that we did not investigate their involvement in other stages. Mast cells colonize peripheral tissues in the fetus ((67), (68), (69), (63)). In the cornea, for example, the first granule-containing mast cells can be found by E12.5 ((70)). It is thus possible that mast cells are already present in the fetal gland, where they could interact with the ductal epithelium. Whilst we did not formally address this, the rudimentary, pre-pubertal tree was indistinguishable between *Karma*^Cre^ Rosa26^lsl-DTA^ mice and mast cell-proficient controls at 3 weeks of age (Figure 2B). Since these mice are devoid of mast cells in the mammary gland at all stages tested, this might indicate that they are also not critical for fetal gland morphogenesis. However, this would need to be investigated specifically at fetal stages. During pregnancy, ducts first undergo additional branching and then later form milk-secreting alveoli (2). Interestingly, the number of mammary gland mast cells increases in mid to late pregnancy, coinciding with alveolar remodeling (71). Furthermore, lactation involves the plasminogen cascade of serine proteases (72) and mast cell granules contain Plasma Kallikrein (PKa), an activator of the cascade (73). Plasminogen and PKa are also needed for involution of the lactating mammary gland, which is triggered by weaning ((72), (73)). This process requires apoptosis of the alveolar epithelium, and repopulation of the mammary fat pad with adipocytes. Mast cells may thus be involved in alveolarization and involution. This could in future work be addressed for example using a DTx-inducible model, as this could avoid maternal mast cell-deficiency during gestation and hence, any potential confounding effects of uterine mast cell deficiency (see above).

Delineating heterogeneity within mammary gland mast cells as well as their potential functional cooperation with other cell types are additional exciting research questions for follow-up work. This could combine single cell omics with genetic approaches to determine their developmental origins and conditionally deplete subsets of mast cells or candidate effectors, such as PKa. *Ms4a2*^lsl-DTR^ will be a valuable tool to disentangle developmental roles for mast cells in the gland, and beyond.

## Acknowledgements

We thank the University of Edinburgh Bioresearch & Veterinary Services for their invaluable services and experimental support, in particular Michael Dodds for outstanding animal husbandry and William Mungall for technical assistance. We further thank the IRR Flow Cytometry Facility. We are also thankful to all other members of the Gentek lab, Calum Bain, Elaine Emmerson and their lab members for feedback and support. We thank Axel Roers for *Mcpt5*-Cre and Pierre Launay for *Ms4a2*^hDTR^ mice.

## Author contributions RG doublecheck when final

Study and experimental design: SK, RG

Conduction, analysis, and/or interpretation of experiments and data: SK, JM, CMM, CC, MMP, BMD

Provision of samples or tools: RA, MD

Technical assistance: SB, GS, KP

Intellectual contribution: GW, AP

Drafting and revision the manuscript: SK, RG

Funding acquisition: RG, SK

## Data and material availability

All data related to this study are available either in the paper or in the Supplementary Materials. Raw data are available upon request.

## Abbreviations

TEBs: Terminal end buds
TEDs: Duct ends

