## Supplementary Figures for "Mast cells are not essential for pubertal mammary gland branching"

### Supplementary Information

**Figure S1 (related to Figure 1):**

**Mast cell abundance in female mammary glands and male fat pads. (A)** Flow cytometric mast cell quantification in the male abdominal fat pad compared to the female mammary gland at adulthood. Data are shown as mean with error bars indicating the SD. *$P < 0.05$ as determined by Mann-Whitney test.

**Figure S1**

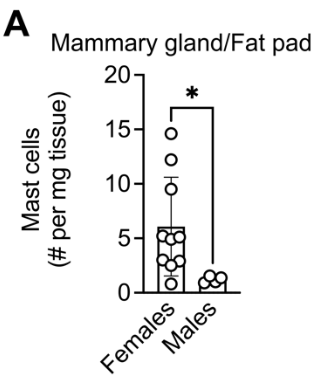


**Supplementary Figure 2 (related to Figure 2): Assessment of mast cell deficiency in other models and at other sites. (A)** Mast cell numbers in the adult mammary glands of *Mcpt5*-Cre:Rosa26$^{lsl\text{-}DTA}$ (red) and control mice (white), indicating lack of targeting. (B) Relative abundance of mast cells in the peritoneal cavity of *Karma*$^{Cre/wt}$ Rosa26$^{lsl\text{-}DTA/wt}$ (red) and *Karma*$^{wt/wt}$ Rosa26$^{lsl\text{-}DTA/wt}$ control (white) at the indicated ages, as measured by flow cytometry. Data are from 46 individual mice from 8 independent experiments and shown as mean with SD as error bars. ***$P < 0.001$, **$P < 0.01$ as determined by Mann-Whitney test. ns = not significant.

**Figure S2**

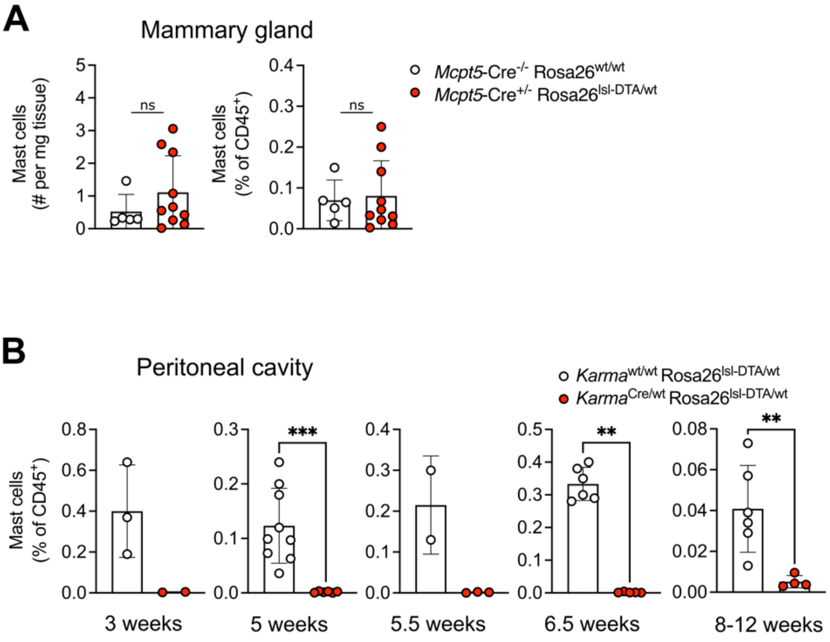


**Supplementary Figure 3 (related to Figure 3): Inducible mast cell depletion by Diphtheria toxin.** (**A, B)** *Ms4a2*$^{hDTR}$ mice were administered Diphtheria toxin (1µg) into the mammary fat pad at 4 weeks and analyzed at puberty (5.5 weeks). Flow cytometry was used to determine mast cell numbers and/or relative abundance within CD45$^{+}$ cells in the mammary gland **(A)** and peritoneal cavity **(B)**.

**Figure S3**

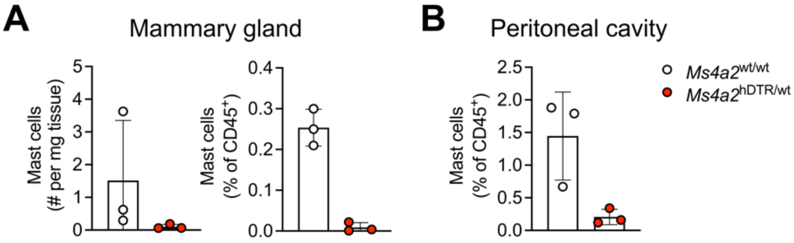
